# *LAMPrey*: a standardised method for analysing quantitative LAMP (qLAMP) and qPCR reactions using the inflection cycle threshold iCt

**DOI:** 10.1101/2025.05.06.651076

**Authors:** Adam Bates, Jiao Li, Francisco Rivero, Katharina C. Wollenberg Valero

## Abstract

Quantitative Loop-mediated isothermal amplification (qLAMP) is a relatively new method that has gained popularity in recent years, particularly in disease identification, including during the recent SARS-CoV-2 pandemic. Unlike conventional quantitative PCR (qPCR), qLAMP features a linear amplification phase before the exponential phase. Determining cycle threshold (Ct) values through automatic thresholding may therefore produce inaccurate results, and the nature of these thresholds complicates comparability between studies and softwares. We introduce a new method for transforming sigmoidal amplification curves into inflection threshold curve (iCt) to address issues with auto thresholds and analysis of qLAMP. This method is implemented as a collection of R functions named LAMPrey, suitable for analysis of both qPCR and qLAMP reactions performed in the two most commonly used real-time thermocyclers. We simulate qLAMP amplification differences, demonstrate that iCt and Ct methods perform equivalently for conventional qPCR with an Illumina library quantitation kit, and show that iCt values outperform Ct values for quantifying qLAMP reactions in zebrafish embryos. All scripts developed for this paper are available at https://github.com/dodged13/LAMPrey

**Graphical Abstract:** 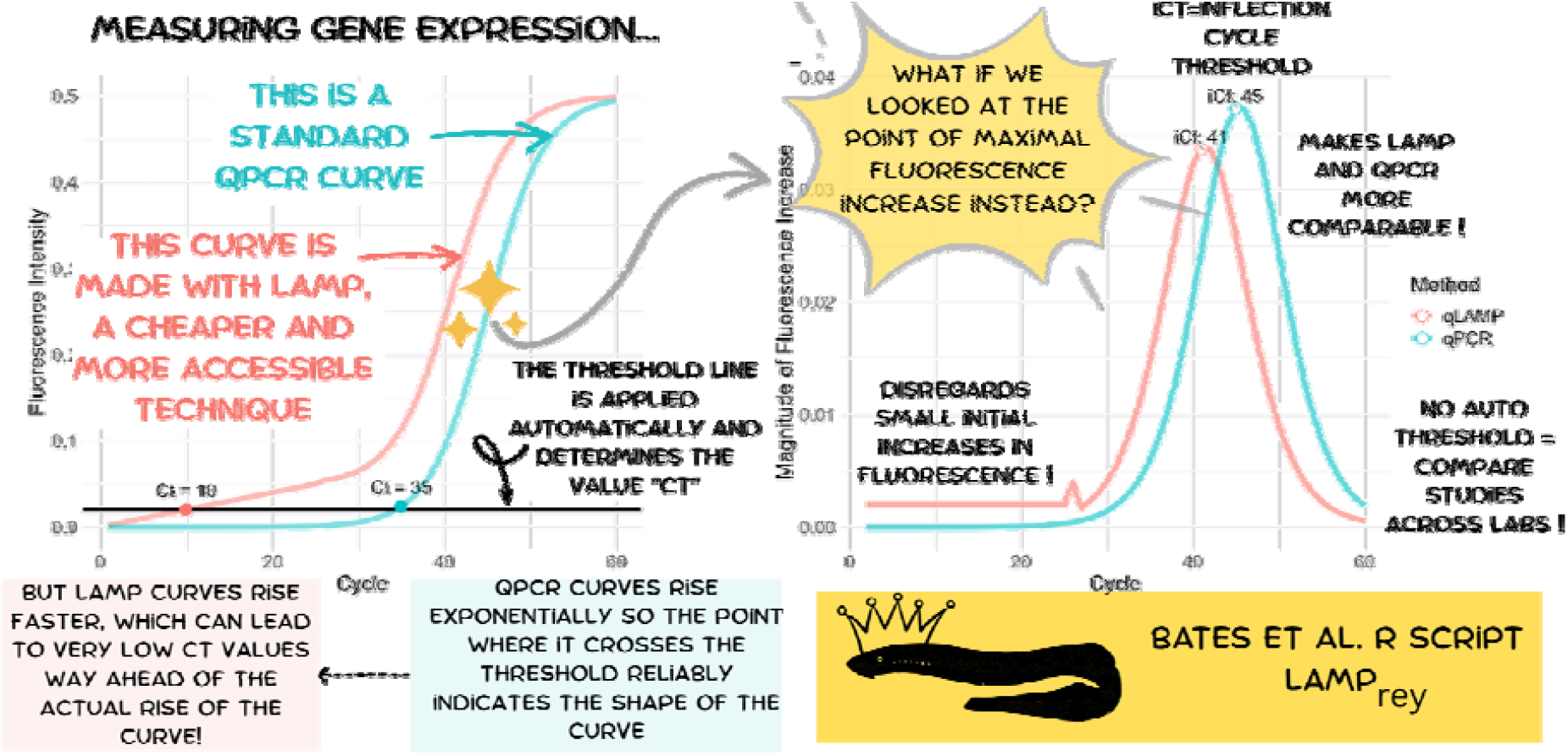

## Introduction

Quantifying specific RNA fragments is a fundamental technique in molecular biology and diagnostics, used to analyse RNA derived from organismal tissue, cells, or infectious agents such as viruses. The current gold standard for RNA quantification is quantitative PCR (qPCR or RT-qPCR), which has the MIQE guidelines in order to standardize the technology and minimise error; which ensures comparability across studies (Bustin et al., 2025). However, qPCR may not be accessible to all labs around the world as it requires a real-time fluorescence detection thermocycler and is also susceptible to non-specific amplification or hairpin formations when using self designed primers (Raso et al, 2011; Bustin & Huggett, 2017). Loop-mediated isothermal amplification PCR (LAMP or qLAMP) offers a lower-cost alternative for gene expression quantification due to its different chemistry and reaction mechanism (Notomi et al, 2000). LAMP gained popularity during the SARS-CoV-2 pandemic for its ease of use in detecting viral presence using pH- or turbidity-dependent reporters and therefore removing the requirement for a real-time thermocycler (Kitagawa et al, 2020; Alves et al, 2021). Unlike conventional PCR, LAMP uses four to six primers that create hairpin structures from the template for continuous amplification that are then extended into high molecular weight structures (“handlebars”). This allows LAMP to be performed on simpler equipment like heating blocks, which lowers the accessibility burden. The technology is also able to use RNA directly as a template, which eliminates the need for reverse transcription, reducing both time and cost and again increasing the accessibility of the technology. The typical LAMP reaction is completed in about 30 minutes, compared to the 45-150 minutes required for qPCR.

In qPCR, manufacturer software implemented in real-time thermal cyclers typically uses an auto-determined or manually set threshold for calculating cycle threshold (Ct) values. Here, proprietary algorithms set a threshold during the exponential phase of amplification, which is applied roughly when the fluorescence of the reaction rises above that of the background noise. This dependency on manufacturer-specific software settings reduces transparency and comparability between studies as Ct values are not standardised between labs or studies. While relative quantification using reference genes is not affected, this lack of standardisation can pose challenges when comparing absolute Ct values, such was observed in the SARS-CoV-2 quantification of viral load (Alves et al, 2021). This approach to Ct calculation also fails to account for variations in reaction efficiency, which can be influenced by contaminants like salts from RNA extraction, leading to inconsistent results between labs. Such issues, combined with inadequate lab experience or training, contributed to the prevalence of false positive cases during the 2019 COVID pandemic (Mouliou & Gourgoulianis, 2021). However, one study found that LAMP had a detection sensitivity of 87% of SARS-CoV-2 positive cases through a colorimetric readout, with 100% specificity making it a practical option for diagnostic labs with limited access to equipment (Lee et al, 2020). For qPCR reactions, the MIQE guidelines on best practice emphasise reproducibility but do not specifically address the impact of threshold settings on comparability across studies (Bustin et al, 2009). In addition, such automatic thresholds can introduce errors when amplification curves deviate from the typical sigmoidal shape of qPCR. qLAMP reactions exhibit an initial linear phase before entering exponential amplification, which can vary further based on the template or primers (Zhang et al, 2023). This variability can lead to erroneous Ct values if the threshold is crossed during the linear phase rather than the exponential phase. Developing transparent and easy to use open-source tools for RNA quantitation will enhance reproducibility and foster open science. Here, we present an alternative method to automatic Ct computation based on the point of maximal fluorescence increase, termed the inflection cycle threshold (iCt). This will eliminate problems with (i) the arbitrary nature of threshold values and (ii) the non-standard initial amplification curves through qLAMP reaction dynamics. This approach enables more consistent quantification across studies, improving the comparability of both qPCR and qLAMP results. We have implemented this method in the R script termed *LAMPrey*.

## Materials and Methods

### Simulation of amplification curves and computing inflection cycle threshold (iCt)

We first simulated typical amplification curves generated by qPCR and qLAMP methods in R (CRAN, R core team, 2024). The qLAMP curve included an initial linear amplification phase, followed by an exponential phase, and concluded with a lag phase when reagents were depleted or all template molecules had been amplified (Appendix 1, https://github.com/dodged13/LAMPrey/blob/main/LAMPreySIM.Rmd).

The *LAMPrey* function was developed as follows: the table of raw fluorescence signals per cycle from a real-time thermal cycler - specifically, the GREEN channel for SYBR green reaction chemistry, though it can be adapted for other channels depending on the dye used - was imported into R. While users can modify the input script to fit the template, it can process raw data files produced by two common real-time thermal cyclers, the StepOnePlus real time thermocycler (Applied Biosystems, Massachusetts) and the QuantStudio real-time PCR system (Applied Biosystems, Massachusetts). As raw fluorescence signals are not standardised, they are first normalised to themselves by dividing each fluorescence reading at cycle n by the reading at cycle n+5, rather than using a background dye like ROX. These normalised values are plotted as a line graph with cycle number on the x-axis. A reactivity curve is generated, with the peak of the curve indicating the cycle where the reaction is most active over 5 cycles (∼90 seconds of LAMP), recorded as the inflection cycle threshold (iCT) for downstream analysis. The *LAMPrey* script is available at (Appendix 2, https://github.com/dodged13/LAMPrey). For ease of use, *LAMPrey* calculates iCT values and returns them for each sample, as well as being able to be used in combination with ggplot2 (Wickham, 2016) to generate publication ready graphs.

### Test of method with qPCR and qLAMP data

iCt values calculated with *LAMPrey* and standard Ct values calculated by the StepOne thermocycler were compared on the example of qPCR data derived from an NEBNext® Illumina library quant kit consisting of a set of standards with known cDNA concentration (New England Biosciences, Massachusetts; Cat #E7630S). qPCR reactions were performed as per manufacturers’ instructions provided with the kit. Subsequently, both machine-based Ct and LAMPrey iCt values were calculated for the same reactions. To compare the performance of Ct and iCt values in determining the threshold of successful amplification for LAMP data, we amplified 14 genes from total RNA of 24 hours post fertilisation zebrafish embryos (*Danio rerio*). Primers were designed using NEB’s LAMP primer design tool, using the .fasta sequences of the genes of interest from NCBI’s RefSeq database and ordered from IDT (Leuven, Belgium). Primers were selected for having closely related melting temperatures (T_M_) as well as having sufficient space between them that loop primers (Loop F and Loop B) could be added to the reaction mix to speed up the reaction. For ease and to reduce pipette errors, each primer set was combined per gene as a 10X stock and added to each reaction as a mix (Table 1). Warmstart LAMP kits were ordered from New England Biosciences (Massachusetts; Cat# E1700).

**Table 1.** The composition and the concentrations of the LAMP primer mix used in the reactions.

| Primer | 10X Concentration | Working concentration |
| --- | --- | --- |
| FIP | 16μM | 1.6μM |
| BIP | 16μM | 1.6μM |
| F3 | 2μM | 0.2μM |
| B3 | 2μM | 0.2μM |
| LOOP F | 4μM | 0.4μM |
| LOOP B | 4μM | 0.4μM |

Across several independent experiments from three researchers (AB, JL, and VS), RNA samples were extracted from pools of 20 zebrafish embryos per each sample. Tissue was first homogenised in Trizol™ using a manual sterile pestle. A standard Trizol™ extraction was performed as per manufacturer’s instructions (Invitrogen, California), although was modified by the addition of an extra ethanol wash step in order to remove phenol contamination. Quality was assessed using a ND-1000 Nanodrop spectrophotometer (Thermofisher, Massachusetts). 70 ng of total RNA has previously been found to be sufficient to generate reliable results from qLAMP (Karthik et al, 2016), and so this was used across all reactions. After initial tests with the 25 uL total reaction volume as suggested by the manufacturer, and a reduced 10 uL reaction volume (Table 2), no differences in effectiveness were detected and 10 uL reactions were subsequently performed. In order to avoid pipetting errors, the fluorescent dye was added to an overall mastermix including primer mix, and LAMP warmstart mastermix. RNA was mixed with water and added in last. Freezing the overall mastermix had no negative effect on reactions. The LAMP reaction itself was performed in a StepOnePlus real time thermocycler (Applied Biosystems, Massachusetts) for 100-200 “cycles” of 18 seconds of which a reading was taken at the end of each 18 second period. With these settings, each reaction has a total reaction time of 30-60 minutes, with temperature set at a constant 65_J. Subsequently, both machine-based Ct and LAMPrey iCt values were calculated for the same reactions, for in total 397 qLAMP reactions for 14 genes. If the Ct and iCt methods produced identical results, a linear regression of one against the other would yield a perfect correlation (R^2^ = 1) with zero residuals. The exponential phase in LAMP and qPCR means that most samples will have both predominantly lower values for Ct and iCt than larger values, and that even very small differences between them represent important biological differences and impact downstream calculations (i.e., ΔCT/ΔiCt and ΔΔCT/ΔΔiCt). Reducing the influence of extreme values through log transformation leads to more stable residuals and a better fit when comparing the variability in iCt relative to Ct. Considering this biological reality, log transformed values of iCt and Ct were compared (script available in Appendix 3 and GitHub).

**Table 2.** Composition of the 10uL qLAMP reactions performed. The reaction can be performed with either RNA or cDNA as the input material.

| Component | Supplier (cat #) | Experimental | No template control |
| --- | --- | --- | --- |
| WarmStart LAMP 2X Master Mix | NEB (E1700S) | 5 μl | 5 μl |
| LAMP Fluorescent Dye (50x) | NEB (B1700SVIAL) | 0.2 μl | 0.2 μl |
| ROX (50x) (optional) | Invitrogen (#12223012) | 0.2 μl | 0.2 μl |
| LAMP Primer Mix (10X) | IDT | 1 μl | 1 μl |
| Target RNA / cDNA | - | 70ng total RNA | - |
| Nuclease free H<sub>2</sub>O | - | Up to 10 μl | Up to 10 μl |

## Results and Discussion

### Current methods are insufficient for qLAMP analysis

Simulated amplification curves for both qPCR and qLAMP chemistry are shown in Figure 1A, illustrating the difference in Ct values when calculated using the same auto threshold. Curves are offset by initially one cycle to avoid overlap. Figure 1B shows the same curves transformed to inflection cycle threshold curves using *LAMPrey*, with iCt being the inflection point of highest value (magnitude of fluorescence increase). This method does not necessitate setting a threshold and, moreover, is not sensitive to the linear phase of qLAMP. The difference between Ct of the two curves equals 25 cycles whereas the difference in iCt using the same data only equals 4 cycles.

**Figure 1.**
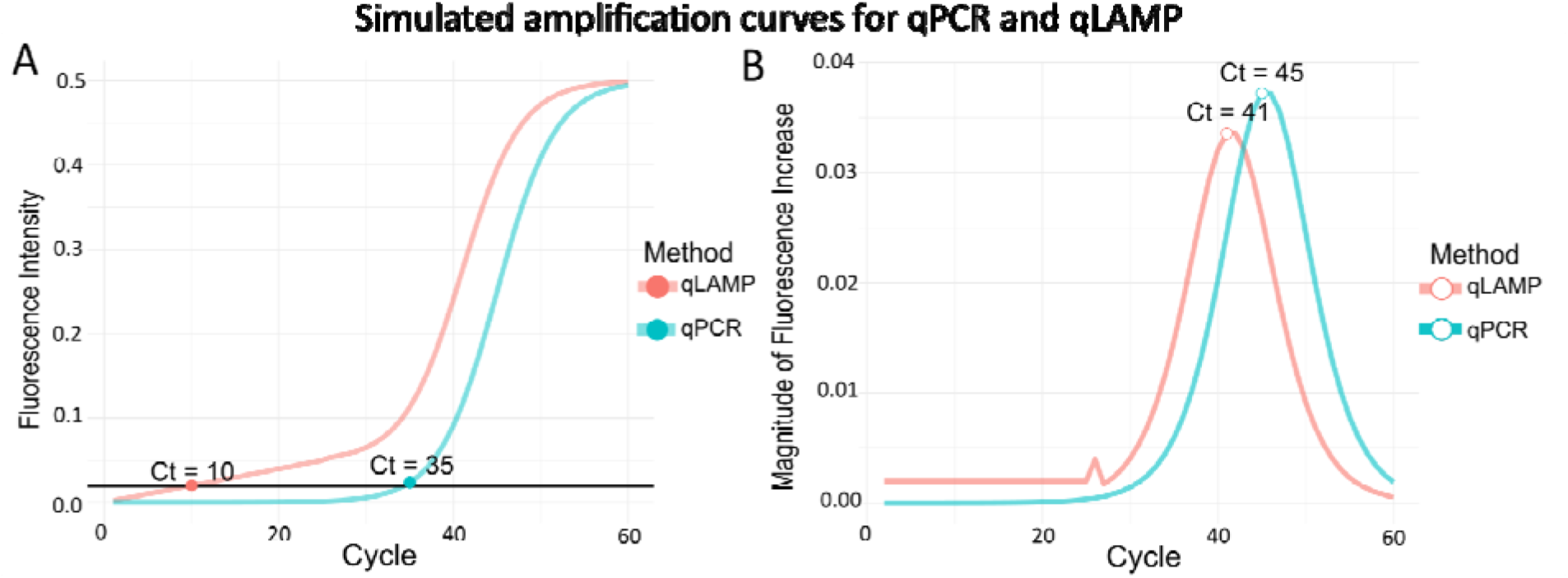
**A)** simulated one-cycle offset qLAMP and qPCR amplification curves, showing the reaction dynamics typical for each assay, namely sigmoidal for qPCR and linear followed by sigmoidal for qLAMP. An automatic threshold is applied to obtain the Ct values, transecting the beginning exponential phase of qPCR, but the linear phase of LAMP, which estimates a lower Ct for qLAMP and a ΔCt=25 between the methods. **B)** The same curves, transformed with LAMPrey to determine the inflection cycle threshold iCt with a difference of ΔiCt=4 between methods.

### LAMPrey iCt can accurately analyse qPCR data

The comparison between the Ct values generated by the StepOne software and the iCt values from the LAMPrey analysis method showed a high correlation, with an R^2^ value of 0.99 (Figure 2). As we used a library quantification kit with given concentrations of DNA standards, we know how much starting DNA is in each reaction. These results show that LAMPrey can accurately detect differences in input concentration of qPCR reactions.

**Figure 2.**
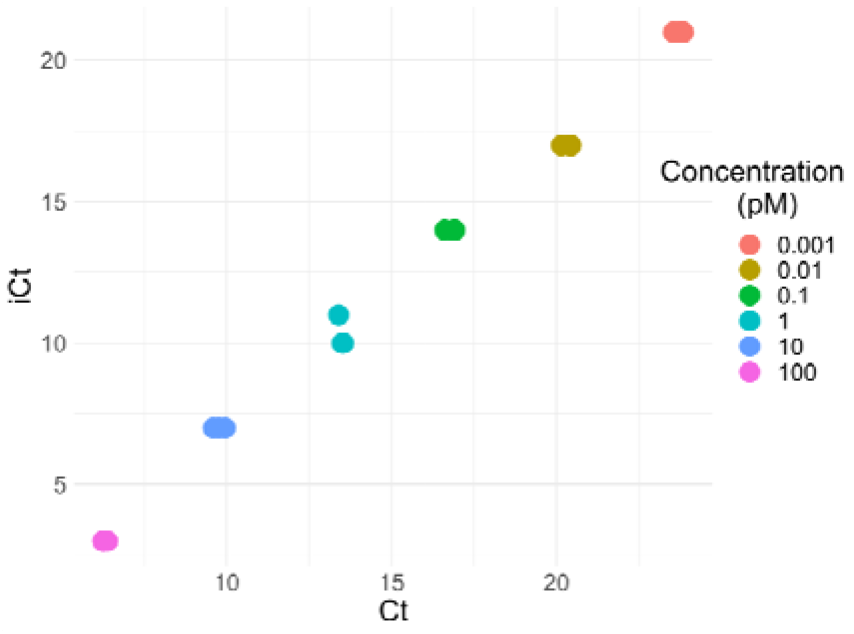
The iCt values calculated for qPCR data of the standards of a NEBNext Quant library kit by the LAMPre method compared to the Ct values provided by the StepOne software. The concentration of the standard is listed as pM. Linear modelling calculated the correlation coefficient (R^2^) between the two methods to be 0.9967

### LAMPrey iCt compared to Ct of zebrafish qLAMP data

The mean of absolute residuals for log(iCt) calculated with *LAMPrey* on log(Ct) calculated with the thermocyclers across all points was 0.0644 while that of log(Ct) on log(iCt) was 0.096. The median of absolute residuals for log(iCt) on logCT across all points was 0.023 while that of log(Ct) on log(iCt) was 0.055. Consequently, log(iCt) values were generally closer to the fit line for both metrics (Figure 3). This suggests that LAMPrey iCt values track the expected amplification dynamics more closely than machine-calculated Ct values in this dataset. Considering small differences and small Ct/iCt values as the optimum of LAMP and qPCR amplification accuracy, this means that iCt slightly outperformed Ct. Notably, residual plots showed that divergence between methods was greatest at low Ct values, suggesting that iCt offers improved precision in cases of high template abundance (=low values), a critical range for biological interpretation.

**Figure 3.**
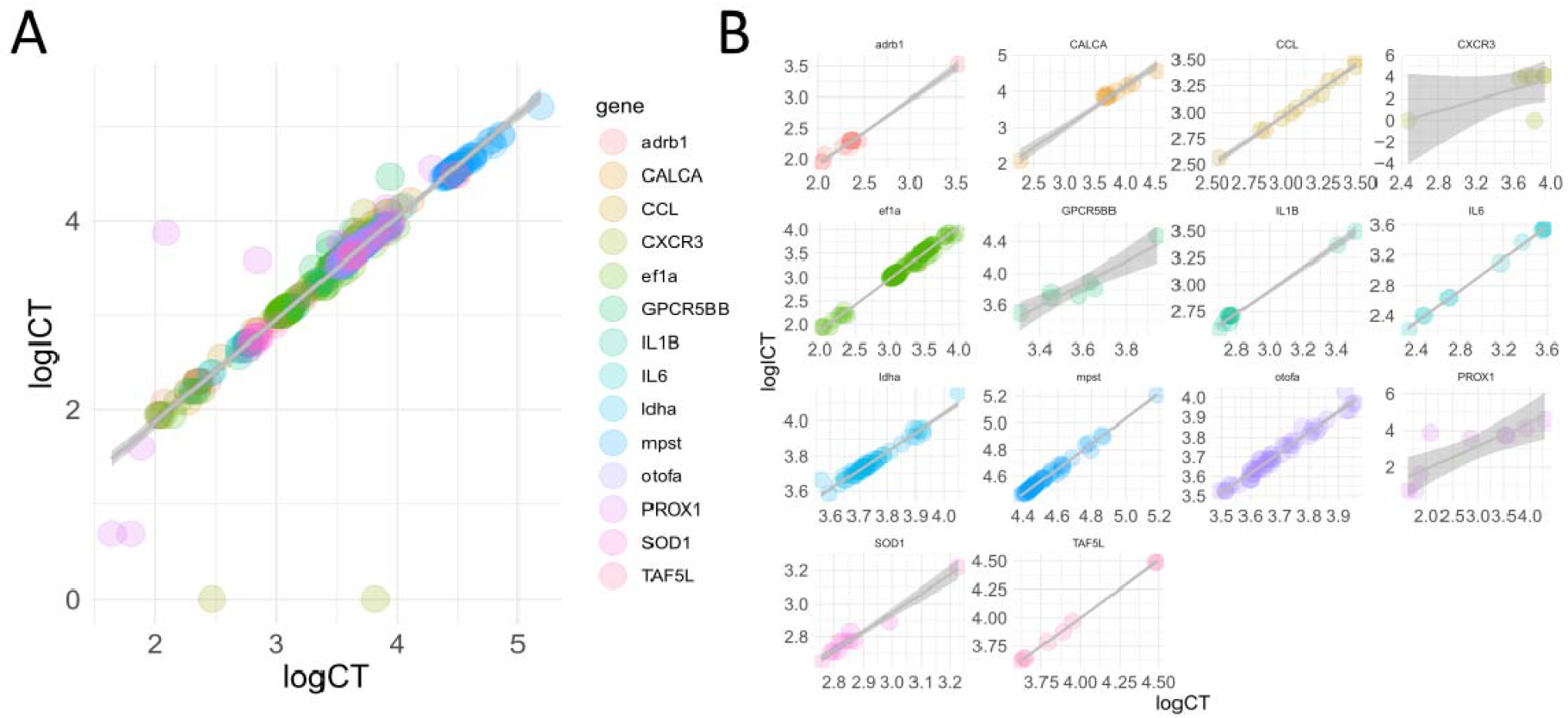
Scatter plot for log(Ct) against log(iCt) from 14 genes amplified with LAMP from zebrafish embryos that were 24 hours post fertilisation (A), and the same scatter plot faceted by gene (B). Linear fit line and 95% confidence intervals are shown in grey.

### LAMPrey can accurately and reliably analyse qLAMP data

As an example, qLAMP amplification curves for the gene *mpst* (mercaptopyruvate sulfurtransferase; seven biological replicates from each n=20 embryos with three technical replicates each) are shown in Figure 4. Mean residual of log(iCt) was 0.009 while that of log(Ct) was 0.0095, which means that for this gene, the *LAMPrey* iCT method also outperforms the Ct method.

**Figure 4.**
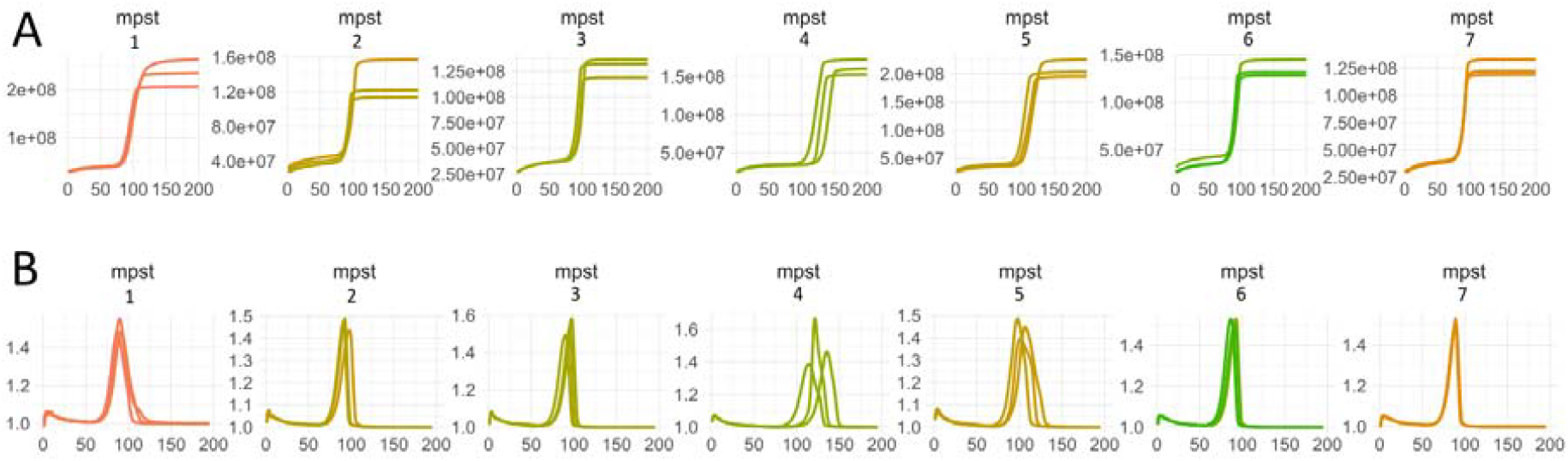
LAMP Amplificati on plots for Ct (A) and iCt (B) for the gene *mpst* which showed lower mean residuals of iCt to the fit line than Ct. The number above each plot is the biological replicate and each line is a technical replicate. The absolute fluorescence plots show a steep rise in the fluorescence in the linear phase, which an auto threshold may count as Ct value, before commencement of the exponential phase. LAMPrey would however not pick these initial rises in fluorescence as the iCt value (small peak in panel B), and use the maximum peak instead.

For ease of use, we have also added a ggplot2 based plotting function called LAMPrey.plot(), which can group the Raw data by desired factor by specifying “well”, “task” or “gene” (Figure 5A, B, C respectively). As the function uses ggplot2, it can be combined with user’s already existing themes, be overlaid with other graphs or annotations and can be faceted to give a better overview of the data (Figure 5D).

**Figure 5.**
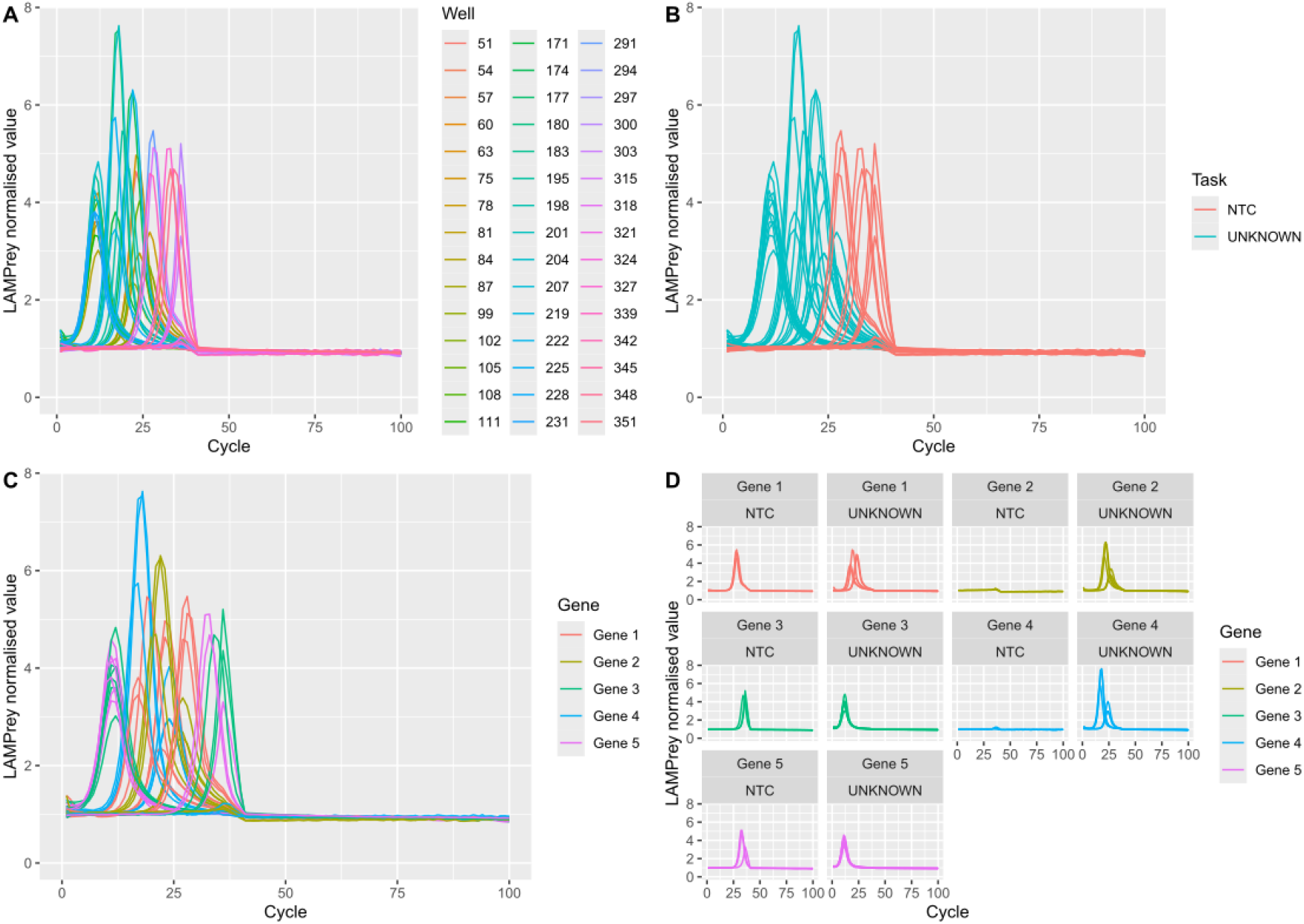
Example outputs of the LAMPrey.plot() function using qPCR data where either **A)** well, **B)** task or **C)** gene has been defined as the grouping variable. **D)** is the gene output but facet wrapped by gene and task (NTC = No Template Control; Unknown = Sample of unknown quantity/Experimental sample) using ggplot’s facet_wrap() function, showing the ability to modify the plot using ggplot functions.

## Conclusion

In conclusion, we have developed an open-source method for RNA quantitation that enhances reproducibility and better supports open science. Our LAMPrey toolkit allows researchers of all abilities to accurately analyse and interpret qLAMP data, thus further reducing the barrier to entry to this already easily accessible technology. Our approach, based on the inflection cycle threshold (iCt), provides a reliable alternative to automatic Ct computation by capturing the point of maximal fluorescence increase independently of thermal cycler or software used. This method addresses the two challenges of arbitrary or proprietary threshold setting and the variable initial amplification dynamics in qLAMP reactions, leading to more consistent and comparable results across studies and technologies. We believe that the LAMPrey method of analysis is a valuable tool for the research community, facilitating improved and reproducible analysis of qPCR and qLAMP data.

## Acknowledgments

We would like to extend our thanks to Victoria Scott (Hull) and Emma Chapman (Hull) for some of the LAMP data, and Graham Sellers (Hull) for providing the NEB library quantification kit data. KWV and JL acknowledge funding by the European Union (ERC, MolStressH2O, #101044202). Views and opinions expressed are however those of the author(s) only and do not necessarily reflect those of the European Union or the European Research Council Executive Agency. Neither the European Union nor the granting authority can be held responsible for them.

## References

Alves, P. A., de Oliveira, E. G., Franco-Luiz, A. P. M., Almeida, L. T., Gonçalves, A. B., Borges, I. A., Rocha, F. de S., Rocha, R. P., Bezerra, M. F., & Miranda, P. (2021). Optimization and clinical validation of colorimetric reverse transcription loop-mediated isothermal amplification, a fast, highly sensitive and specific COVID-19 molecular diagnostic tool that is robust to detect SARS-CoV-2 variants of concern. Frontiers in Microbiology, 12, 713713.

Broeders, S., Huber, I., Grohmann, L., Berben, G., Taverniers, I., Mazzara, M., Roosens, N., & Morisset, D. (2014). Guidelines for validation of qualitative real-time PCR methods. Trends in Food Science & Technology, 37(2), 115–126.

Bustin, S. A., Ruijter, J. M., van den Hoff, M. J. B., Kubista, M., Pfaffl, M. W., Shipley, G. L., Tran, N., Rödiger, S., Untergasser, A., Mueller, R., Nolan, T., Milavec, M., Burns, M. J., Huggett, J. F., Vandesompele, J., & Wittwer, C. T. (2025). MIQE 2.0: Revision of the Minimum Information for Publication of Quantitative Real-Time PCR Experiments Guidelines. Clinical Chemistry, hvaf043.

Bustin, S., & Huggett, J. (2017). qPCR primer design revisited. Biomolecular Detection and Quantification, 14, 19–28.

Huggett, J., Dheda, K., Bustin, S., & Zumla, A. (2005). Real-time RT-PCR normalisation; strategies and considerations. Genes & Immunity, 6(4), 279–284.

Karthik, K., Rathore, R., Thomas, P., Viswas, K. N., Agarwal, R. K., Rekha, V., Jagapur, R. V., & Dhama, K. (2016). Rapid and visual loop mediated isothermal amplification (LAMP) test for the detection of Brucella spp. And its applicability in epidemiology of bovine brucellosis. Veterinarski Arhiv, 86(1), 35–47.

Kitagawa, Y., Orihara, Y., Kawamura, R., Imai, K., Sakai, J., Tarumoto, N., Matsuoka, M., Takeuchi, S., Maesaki, S., & Maeda, T. (2020a). Evaluation of rapid diagnosis of novel coronavirus disease (COVID-19) using loop-mediated isothermal amplification. Journal of Clinical Virology, 129, 104446.

Kitagawa, Y., Orihara, Y., Kawamura, R., Imai, K., Sakai, J., Tarumoto, N., Matsuoka, M., Takeuchi, S., Maesaki, S., & Maeda, T. (2020b). Evaluation of rapid diagnosis of novel coronavirus disease (COVID-19) using loop-mediated isothermal amplification. Journal of Clinical Virology, 129, 104446.

Lee, J. Y., Best, N., McAuley, J., Porter, J. L., Seemann, T., Schultz, M. B., Sait, M., Orlando, N., Mercoulia, K., & Ballard, S. A. (2020). Validation of a single-step, single-tube reverse transcription loop-mediated isothermal amplification assay for rapid detection of SARS-CoV-2 RNA. Journal of Medical Microbiology, 69(9), 1169–1178.

Mouliou, D. S., & Gourgoulianis, K. I. (2021). False-positive and false-negative COVID-19 cases: Respiratory prevention and management strategies, vaccination, and further perspectives. Expert Review of Respiratory Medicine, 15(8), 993–1002.

Notomi, T., Okayama, H., Masubuchi, H., Yonekawa, T., Watanabe, K., Amino, N., & Hase, T. (2000). Loop-mediated isothermal amplification of DNA. Nucleic Acids Research, 28(12), e63– e63.

R Core Team. (2024). R: A Language and Environment for Statistical Computing. R Foundation for Statistical Computing. https://www.R-project.org/

Raso, A., Mascelli, S., Nozza, P., Ugolotti, E., Vanni, I., Capra, V., & Biassoni, R. (2011). Troubleshooting fineLJtuning procedures for qPCR system design. Journal of Clinical Laboratory Analysis, 25(6), 389–394.

Schefe, J. H., Lehmann, K. E., Buschmann, I. R., Unger, T., & Funke-Kaiser, H. (2006). Quantitative real-time RT-PCR data analysis: Current concepts and the novel “gene expression’s CT difference” formula. Journal of Molecular Medicine, 84, 901–910.

Wickham, H. (2016). ggplot2: Elegant Graphics for Data Analysis. Springer-Verlag New York. https://ggplot2.tidyverse.org

Zhang, X., Zhao, Y., Zeng, Y., & Zhang, C. (2023). Evolution of the probe-based loop-mediated isothermal amplification (LAMP) assays in pathogen detection. Diagnostics, 13(9), 1530.

