## Supplementary materials for "*LAMPrey*: a standardised method for analysing quantitative LAMP (qLAMP) and qPCR reactions using the inflection cycle threshold iCt"

**Appendix**

**Appendix 1**: *R Markdown Raw File used for the generation of the simulation curves in figure 1 also available at https://github.com/dodged13/LAMPrey*

---

title: "LAMPrey data simulation"

author: "Kat Valero"

date: "`r Sys.Date()`"

output: html_document

---

```{r setup, include=FALSE}

knitr::opts_chunk$set(echo = TRUE)

```

\## R Markdown

This is an R Markdown document. Markdown is a simple formatting syntax for authoring HTML, PDF, and MS Word documents. For more details on using R Markdown see <http://rmarkdown.rstudio.com>.

When you click the **Knit** button a document will be generated that includes both content as well as the output of any embedded R code chunks within the document. You can embed an R code chunk like this:

```{r cars}

summary(cars)

```

```{r}

library(tidyverse)

\# Set parameters

set.seed(123) # For reproducibility

n_cycles <- 60 # Number of cycles

initial_intensity <- 1e-4 # Initial fluorescence intensity

max_intensity <- 0.5 # Maximum fluorescence intensity for both reactions

\# Sigmoidal growth function

sigmoidal_growth <- function(cycle, start_intensity, max_intensity, midpoint, rate) {

start_intensity + (max_intensity - start_intensity) / (1 + exp(-rate * (cycle - midpoint)))

}

\# Function to simulate qPCR data (sigmoidal growth)

simulate_qpcr <- function(n_cycles, initial_intensity, max_intensity) {

cycles <- 1:n_cycles

midpoint <- 45 # Midpoint for the sigmoidal growth

rate <- 0.3 # Rate for the sigmoidal growth

intensity_qpcr <- sapply(cycles, function(cycle) sigmoidal_growth(cycle, initial_intensity, max_intensity, midpoint, rate))

return(data.frame(Cycle = cycles, Intensity = intensity_qpcr, Method = "qPCR"))

}

\# Function to simulate qLAMP data (initial linear growth followed by sigmoidal growth)

simulate_qlamp <- function(n_cycles, initial_intensity, max_intensity) {

cycles <- 1:n_cycles

linear_phase <- 25 # Length of the initial linear phase

linear_rate <- 0.002 # Rate of increase for the linear phase

sigmoid_rate <- 0.3 # Rate for the sigmoidal growth

midpoint <- 13 # Midpoint for the sigmoidal growth

intensity_qlamp <- numeric(n_cycles)

for (i in 1:n_cycles) {

if (i <= linear_phase) { # Initial linear phase

intensity_qlamp[i] <- initial_intensity + i * linear_rate

} else { # Sigmoidal growth phase

start_intensity <- (initial_intensity - 0.001) + linear_phase * linear_rate

intensity_qlamp[i] <- sigmoidal_growth(i - linear_phase - 3, start_intensity, max_intensity, midpoint, sigmoid_rate)

}

}

return(data.frame(Cycle = cycles, Intensity = intensity_qlamp, Method = "qLAMP"))

}

\# Simulate data

qpcr_data <- simulate_qpcr(n_cycles, initial_intensity, max_intensity)

qlamp_data <- simulate_qlamp(n_cycles, initial_intensity, max_intensity)

\# Combine data

simulated_data <- bind_rows(qpcr_data, qlamp_data)

\# Define the threshold

threshold <- 0.02

\# Find the intersection points

intersection_points <- simulated_data %>%

group_by(Method) %>%

filter(Intensity >= threshold) %>%

slice(1) %>% # Take the first point where intensity crosses the threshold

ungroup()

\# Plot the simulated data with the threshold line and labeled intersection points

plotCT <- ggplot(simulated_data, aes(x = Cycle, y = Intensity, color = Method)) +

geom_line(size = 2, alpha=0.5) +

geom_hline(yintercept = threshold, size= 1.5, color = "black") + # Add the threshold line

geom_point(data = intersection_points, aes(x = Cycle, y = Intensity), size = 4) +

geom_text(data = intersection_points, aes(x = Cycle, y = Intensity, label = paste0("Ct = ", Cycle)),

vjust = -1, hjust = 0.5, size = 5, color = "black") +

labs(title =" ",

x = "Cycle", y = "Fluorescence Intensity") +

theme_minimal()

```

Calculate the LAMPrey method for the simulated data

```{r}

\# Load necessary libraries

library(tidyverse)

library(ggrepel)

\# Read the simulated data

simulated_data <- read.csv("simulated_amplification_data.csv")

\# Calculate the magnitude of fluorescence increase between each cycle

simulated_data <- simulated_data %>%

group_by(Method) %>%

mutate(Magnitude_Increase = Intensity - lag(Intensity, default = first(Intensity)))

\# Remove the first row of each group to avoid the initial zero increase

simulated_data <- simulated_data %>%

filter(row_number() != 1)

\# Identify the cycle with the highest increase for each method

max_increase <- simulated_data %>%

group_by(Method) %>%

slice(which.max(Magnitude_Increase))

\# Plot the magnitude of fluorescence increase per step for each method

plotICT<- ggplot(simulated_data, aes(x = Cycle, y = Magnitude_Increase, color = Method)) +

geom_line(size = 2, alpha=0.6) +

geom_point(data = max_increase, aes(x = Cycle, y = Magnitude_Increase), size = 4, shape = 21, fill = "white") +

geom_text_repel(data = max_increase, aes(x = Cycle, y = Magnitude_Increase, label = paste("iCt:", Cycle)),

nudge_y = 0.001, size = 5, color = "black",

segment.color = "grey50", min.segment.length = 0.1) +

scale_y_continuous(limits = c(0, NA)) + # Extend the Y-axis to ensure labels are visible

labs(title = " ", x = "Cycle", y = "Magnitude of Fluorescence Increase") +

theme_minimal()

```

```{r}

\# Save plotCT as a PDF

ggsave("plotCT.pdf", plot = plotCT, width = 8, height = 6)

\# Save plotICT as a PDF

ggsave

\# Increase font sizes in plotCT

plotCT <- plotCT +

theme(

plot.title = element_text(size = 20, face = "bold"),

axis.title = element_text(size = 16),

axis.text = element_text(size = 14),

legend.title = element_text(size = 16),

legend.text = element_text(size = 14)

)

\# Increase font sizes in plotICT

plotICT <- plotICT +

theme(

plot.title = element_text(size = 20, face = "bold"),

axis.title = element_text(size = 16),

axis.text = element_text(size = 14),

legend.title = element_text(size = 16),

legend.text = element_text(size = 14)

)

#install.packages("cowplot")

library(cowplot)

\# Combine the plots with labels "A" and "B"

combined_plot <- plot_grid(

plotCT + ggtitle("A") + theme(plot.title = element_text(hjust = -0.1, size = 16)),

plotICT + ggtitle("B") + theme(plot.title = element_text(hjust = -0.1, size = 16)),

labels = NULL, # No need for automatic labels since we're using titles

ncol = 2 # Place them side by side

)

\# Save the combined plot as a PDF

ggsave("combined_plot.pdf", plot = combined_plot, width = 16, height = 6)

\# Save the combined plot as a PDF

ggsave("combined_plot.png", plot = combined_plot, width = 16, height = 6)

```

Simulate a curve for the addition of ROX dye

```{r}

\# Load necessary libraries

\# Load necessary libraries

library(ggplot2)

\# Simulate cycle numbers

cycles <- 1:40

\# Parameters for the simulation

normal_volume_baseline <- 100 # Baseline fluorescence for normal volume

higher_volume_baseline <- 150 # Higher baseline due to increased volume

\# Sigmoidal curve parameters

efficiency <- 1.9

amplification_start <- 15

plateau <- 300 # Plateau phase fluorescence

\# Sigmoidal amplification curve adjusted for different baselines

amplification_curve <- function(cycle, baseline, efficiency, start, plateau) {

baseline + (plateau - baseline) / (1 + exp(-0.3 * (cycle - start)))

}

\# Generate the amplification curves

normal_volume_curve <- amplification_curve(cycles, normal_volume_baseline, efficiency, amplification_start, plateau)

higher_volume_curve <- amplification_curve(cycles, higher_volume_baseline, efficiency, amplification_start, plateau)

\# Create a data frame for plotting

data <- data.frame(

Cycle = rep(cycles, 2),

Fluorescence = c(normal_volume_curve, higher_volume_curve),

Volume = rep(c("Normal Volume", "Higher Volume"), each = length(cycles))

)

\# Plotting the curves using ggplot2

ggplot(data, aes(x = Cycle, y = Fluorescence, color = Volume)) +

geom_line(size = 1.2) +

geom_hline(yintercept = 200, linetype = "dashed", color = "gray") +

scale_y_continuous(limits = c(0, 400)) +

scale_x_continuous(limits = c(1, 40)) +

labs(title = "Adjusted Sigmoidal qPCR Curves Without ROX Dye Normalization",

x = "Cycle",

y = "Fluorescence") +

theme_minimal() +

theme(legend.title = element_blank())

```

**Appendix 2:** *The LAMPrey R functions, also available at https://github.com/dodged13/LAMPrey*

LAMPrey.Read_StepOne = function(){

library(readxl)

x = file.choose()

Setup <<- data.frame(read_excel(x, sheet = "Results"))

z = which(Setup == "Task",arr.ind = TRUE)[1]

colnames(Setup) <<- Setup[z,]

Setup <<- Setup[z+1:nrow(Setup),]

Setup <<- data.frame(read_excel(x, sheet = "Results", skip = z))

Raw_Data <<- data.frame(read_excel(x, sheet = "Raw Data", skip = 6))

}

\### Reads in the data from an excel output from a StepOne reaction, specifically raw data and results ###

LAMPrey.Read_QuantStudio = function(){

library(readxl)

x = file.choose()

Setup <<- data.frame(read_excel(x, sheet = "Sample Setup"))

z = which(Setup == "Task",arr.ind = TRUE)[1]

colnames(Setup) <<- Setup[z,]

Setup <<- Setup[z+1:nrow(Setup),]

Raw_Data <<- data.frame(read_excel(x, sheet = "Raw Data", skip = z))

}

\### Reads in the data from an excel output from a QuantStudio reaction, specifically raw data and results ###

LAMPrey.annotate_StepOne = function() {

Raw_Data$Gene <<- Setup$Target.Name[pmatch(Raw_Data$Well, Setup$Well, duplicates.ok = TRUE)]

Raw_Data$Task <<- Setup$Task[pmatch(Raw_Data$Well, Setup$Well, duplicates.ok = TRUE)]

}

LAMPrey.annotate_QuantStudio = function() {

Raw_Data$Gene <<- Setup$'Target Name'[pmatch(Raw_Data$Well, Setup$Well, duplicates.ok = TRUE)]

Raw_Data$Task <<- Setup$Task[pmatch(Raw_Data$Well, Setup$Well, duplicates.ok = TRUE)]

} ### annotates the raw data with columns detailing the genes and the task of the well (standard/unknown) ###

LAMPrey.normalise = function(x) {

well_list = unique(na.omit(Raw_Data)$Well)

cycle_list = unique(na.omit(Raw_Data)$Cycle)

for (i in unique(well_list)) {

for (cy in cycle_list) {

Raw_Data$normalised[Raw_Data$Well == i & Raw_Data$Cycle == cycle_list[cy]] <<- Raw_Data[[x]][Raw_Data$Well == i & Raw_Data$Cycle == cycle_list[cy+5]] /

(Raw_Data[[x]][Raw_Data$Cycle == cycle_list[cy] & Raw_Data$Well == i ])

} }

return(Raw_Data)} ### Uses the colour channel name as input - x = "GREEN" for Step one or "x1.m1" for QuantStudio, to calculate the Normalised LAMP value ###

LAMPrey.plot = function(x) {

library(ggplot2)

if(missing(x)){ print("Please specify 'Task', 'Gene' or 'Well' as a grouping factor")

}else if(x == "Well" | x == "well") { ggplot(data = Raw_Data[!is.na(Raw_Data$Gene),]) +

geom_line(aes(x = Cycle, y = normalised, colour = as.factor(Well)), size = 0.5) +

xlim(0,100) +

labs(y = "LAMPrey normalised value", colour = "Well")

}else if(x == "Task" | x =="task") {

ggplot(data = Raw_Data[!is.na(Raw_Data$Gene),]) +

geom_line(aes(x = Cycle, y = normalised, group = as.factor(Well), colour = as.factor(Task)), size = 0.5) +

xlim(0,100) +

labs(y = "LAMPrey normalised value", colour = "Task")

}else if(x == "Gene" | x =="gene") {

ggplot(data = Raw_Data[!is.na(Raw_Data$Gene),]) +

geom_line(aes(x = Cycle, y = normalised, group = as.factor(Well), colour = as.factor(Gene)), size = 0.5) +

xlim(0,100) +

labs(y = "LAMPrey normalised value", colour = "Gene")

}

}

LAMPrey.analyse = function() {

well_list = unique(na.omit(Raw_Data)$Well)

LAMPrey_Results <<- data.frame(Well = well_list)

for (i in unique(well_list)) {

LAMPrey_Results$Gene_stepone[LAMPrey_Results$Well == i] <<- Setup$Target.Name[Setup$Well == i]

LAMPrey_Results$Gene_quantstudio[LAMPrey_Results$Well == i] <<- Setup$'Target Name'[Setup$Well == i]

LAMPrey_Results$Task[LAMPrey_Results$Well == i] <<- Setup$Task[Setup$Well == i]

LAMPrey_Results$LAMPrey[LAMPrey_Results$Well == i] <<- approx(Raw_Data$normalised[Raw_Data$Well == i],

Raw_Data$Cycle[Raw_Data$Well == i],

max(na.omit(Raw_Data$normalised[Raw_Data$Well == i])))[2] }

} ### Calculates a CT approximation for each well and creates a result dataframe named LAMPrey_Results ###

**Appendix 3:** *The code used to generate a linear model of Ct vs iCt data, also available at https://github.com/dodged13/LAMPrey*

\# Starting with a csv file containing CT and ICT values for each gene

\# Step 1: Fit the linear model ICT ~ CT

lm_ict_on_ct <- lm(log(ICT) ~ log(CT), data = data)

data$residuals_ict_on_ct <- abs(resid(lm_ict_on_ct))

\# Step 2: Fit the linear model CT ~ ICT (inverted model)

lm_ct_on_ict <- lm(log(CT) ~ log(ICT), data = data)

data$residuals_ct_on_ict <- abs(resid(lm_ct_on_ict))

\# Step 3: Calculate and compare the mean absolute residuals

mean_residuals_ict_on_ct <- mean(data$residuals_ict_on_ct)

mean_residuals_ct_on_ict <- mean(data$residuals_ct_on_ict)

print(paste("Mean absolute residuals for logICT on logCT:", mean_residuals_ict_on_ct))

print(paste("Mean absolute residuals for logCT on logICT:", mean_residuals_ct_on_ict))

\# Determine which variable is closer to the linear fit

if (mean_residuals_ict_on_ct < mean_residuals_ct_on_ict) {

print("logICT values are generally closer to the linear fit line.")

} else if (mean_residuals_ict_on_ct > mean_residuals_ct_on_ict) {

print("logCT values are generally closer to the linear fit line.")

} else {

print("logCT and logICT values are equally close to the linear fit line.")

}

\# Step 3b: Calculate and compare the median absolute residuals

median_residuals_ict_on_ct <- median(data$residuals_ict_on_ct)

median_residuals_ct_on_ict <- median(data$residuals_ct_on_ict)

print(paste("Median absolute residuals for logICT on logCT:", median_residuals_ict_on_ct))

print(paste("Median absolute residuals for logCT on logICT:", median_residuals_ct_on_ict))

\# Determine which variable is closer to the linear fit

if (median_residuals_ict_on_ct < median_residuals_ct_on_ict) {

print("logICT values are generally closer to the linear fit line.")

} else if (median_residuals_ict_on_ct > median_residuals_ct_on_ict) {

print("logCT values are generally closer to the linear fit line.")

} else {

print("logCT and logICT values are equally close to the linear fit line.")

}

library(dplyr)

\# Group the data by gene2 and perform calculations for each group

results <- data %>%

group_by(gene) %>%

summarise(

mean_residuals_ict_on_ct = mean(abs(resid(lm(log(ICT) ~ log(CT), data = cur_data())))),

mean_residuals_ct_on_ict = mean(abs(resid(lm(log(CT) ~ log(ICT), data = cur_data()))))

) %>%

mutate(

closer_to_fit = case_when(

mean_residuals_ict_on_ct < mean_residuals_ct_on_ict ~ "ICT",

mean_residuals_ict_on_ct > mean_residuals_ct_on_ict ~ "CT",

TRUE ~ "Equal"

)

)

\# View the results

print(results)

\# Save the results to a CSV file (optional)

write.csv(results, "logCT_vs_logICT_closeness_by_gene.csv", row.names=FALSE)
